# Metabolomic and transcriptomic remodeling of bone marrow myeloid cells in response to maternal obesity

**DOI:** 10.1101/2024.08.20.608809

**Authors:** Yem J Alharithi, Elysse A. Phillips, Tim D. Wilson, Sneha P. Couvillion, Carrie D. Nicora, Priscila Darakjian, Shauna Rakshe, Suzanne S. Fei, Brittany Counts, Thomas O. Metz, Robert Searles, Sushil Kumar, Alina Maloyan

## Abstract

Maternal obesity puts the offspring at high risk of developing obesity and cardio-metabolic diseases in adulthood. Here, using a mouse model of maternal high-fat diet (HFD)-induced obesity, we show that whole body fat content of the offspring of HFD-fed mothers (Off-HFD) increases significantly from very early age when compared to the offspring regular diet-fed mothers (Off-RD). We have previously shown significant metabolic and immune perturbations in the bone marrow of newly-weaned offspring of obese mothers. Therefore, we hypothesized that lipid metabolism is altered in the bone marrow Off-HFD in newly-weaned offspring of obese mothers when compared to the Off-RD. To test this hypothesis, we investigated the lipidomic profile of bone marrow cells collected from three-week-old offspring of regular and high fat diet-fed mothers. Diacylgycerols (DAGs), triacylglycerols (TAGs), sphingolipids and phospholipids, including plasmalogen, and lysophospholipids were remarkably different between the groups, independent of fetal sex. Levels of cholesteryl esters were significantly decreased in offspring of obese mothers, suggesting reduced delivery of cholesterol to bone marrow cells. This was accompanied by age-dependent progression of mitochondrial dysfunction in bone marrow cells. We subsequently isolated CD11b+ myeloid cells from three-week-old mice and conducted metabolomics, lipidomics, and transcriptomics analyses. The lipidomic profiles of these bone marrow myeloid cells were largely similar to that seen in bone marrow cells and included increases in DAGs and phospholipids alongside decreased TAGs, except for long-chain TAGs, which were significantly increased. Our data also revealed significant sex-dependent changes in amino acids and metabolites related to energy metabolism. Transcriptomic analysis revealed altered expression of genes related to major immune pathways including macrophage alternative activation, B-cell receptor signaling, TGFβ signaling, and communication between the innate and adaptive immune systems. All told, this study revealed lipidomic, metabolomic, and gene expression abnormalities in bone marrow cells broadly, and in bone marrow myeloid cells particularly, in the newly-weaned offspring of obese mothers, which might at least partially explain the progression of metabolic and cardiovascular diseases in their adulthood.

## INTRODUCTION

Obesity and metabolic diseases are the biggest epidemics in history and major challenges to healthcare systems worldwide (1). While family history plays an important role, only 20% of cases are explained by genetic predisposition (2), which is often accompanied by a so-called “second hit”, which can include effects from lifestyle choices regarding diet, physical activity, and medication with obesogenic side effects (3). However, it is now widely established that prenatal exposure to an adverse maternal environment, such as maternal obesity, increases the risk of obesity and associated cardio-metabolic diseases in a phenomenon called developmental or fetal programming (4).

In the US, the prevalence of overweight and obesity among women of reproductive age (15-49 years old) has increased from 28.4% in 1999-2000 to 41.5% in 2016 (5). By extension, obesity in women of reproductive age is becoming a serious public health issue; it is associated with reduced fertility, increased risk of pregnancy complications, and long-term health complications for both mother and offspring (6). In particular, obesity in pregnancy has been associated with major birth defects, including neural tube and cardiac defects (7), and with increased risk of later-life neurodevelopmental disorders, including cerebral palsy, attention- deficit disorder, cognitive delay, and even autism (8–10). Importantly, children born to obese mothers are themselves at much higher risk of becoming obese and developing cardiovascular morbidity or type 2 diabetes (11). Published data have revealed that an adverse metabolic environment affects bone marrow activation, leading to cardio-metabolic diseases and increased inflammation. The bone marrow is the primary site of hematopoiesis, and various physiological and pathological characteristics of the marrow regulate hematopoietic progenitor proliferation and migration through blood into tissues (12). Recently, altered bone marrow metabolism has been linked to early signs of metabolic syndrome and atherogenesis in humans (13), and glucose metabolism in the marrow has been shown to correlate with severity of systemic inflammation (14). Obesity is associated with dysregulations in bone marrow homeostasis (15), and a mouse model of diet-induced obesity exhibits myeloid cell expansion and increased production of pro-inflammatory macrophages (16) along with metabolic reprogramming of myeloid cells in response to changing metabolic conditions (17). However, the effects of maternal obesity on the offspring’s myeloid cells and metabolism have not yet been fully examined.

To understand the effects of maternal obesity on immune system development in the offspring, we previously conducted a study on the immune complexity and energy metabolism of bone marrow cells (18). Although that study provided many important insights, the effect of maternal obesity on lipid mediators in offspring bone marrow remained unexamined.

The present work aimed to close this gap by conducting lipidomics analyses of bone marrow cells. Additionally, in order to understand the effects of maternal obesity on offspring innate immunity, we conducted metabolomics and transcriptomics analyses of bone marrow myeloid cells. Our findings reveal distinct, coordinated shifts of lipid mediator profiles and gene expression in bone marrow myeloid cells in response to maternal obesity.

## MATERIALS AND METHODS

### Materials

The EasySep™ Mouse CD11b Positive Selection Kit II was purchased from STEMCELL Technologies (Cambridge, MA).

### Study approval

All animal experiments were approved by the Oregon Health & Science University’s Institutional Animal Use Committee (Protocol # IP00432).

### Mouse model of maternal obesity and ethical statement

All studies were performed using wild-type FVB/N mice. Mice were kept under a 12-hour light/dark cycle, between 18-23 °C with 40-60% humidity, in stress-free/bacteria-free conditions. Mice were caged in groups of three to five whenever possible. Food and water were given ad libitum. Body weights were collected weekly. In this investigation of diet-induced maternal obesity, a high-fat diet (HFD, Teklad Cat#TD.06415, Inotiv, West Lafayette, IN) or the control regular diet (RD, PicoLab® Laboratory Rodent Diet Cat #5L0D, LabDiet, St. Louis, Missouri) was given to virgin female FVB/NJ mice from six weeks of age and throughout the entire study. The composition of these diets has been reported previously (19). After eight weeks of dietary intervention, RD- and HFD-fed female mice were bred to an age-matched RD-fed male. Breeding couples were fed based on the female’s pre-pregnancy diet. At weaning, male and female offspring from each mother were randomly selected for study, with offspring groups designated according to maternal diet as Off-RD and Off-HFD respectively. Both male and female offspring were included in this study, and analyses were performed using disaggregated data. If no differences were observed between the sexes, the data were combined. Every mouse in each experimental group belonged to a different litter. Offspring were fed the regular diet only, starting from weaning and throughout their life.

### Body fat mass quantitation using EchoMRI

Body composition was measured using an EchoMRI Analyser E26-206-RMT (EchoMRI LLC, Houston, TX), as we have previously described (19). The measured parameters included fat and lean mass, and free and total water. Total water encompasses free water and lean mass water. Outcomes are made relative to body mass at timepoint.

### Isolation of bone marrow cells

Bone marrow cells were isolated as previously reported (20). Briefly, femurs were flushed using a 25G5/8 needle with medium containing RPMI1640 + 2% FBS + 10 units/ml heparin + penicillin and streptomycin. Bits of bone were removed using a sterile 4 mm nylon cell strainer (Falcon 352340). Cells were dissolved in 50 ml medium and centrifuged at 2000 rpm (900 x *g*) for ten minutes at 4 °C. The resultant cell pellets were washed twice with 50 ml of serum-free RPMI (RPMI1640 + 20 mM Hepes + penicillin and streptomycin, adjusted to pH 7.4 before filtering), centrifuged at 2000 rpm at 4 °C for five minutes, and finally resuspended in 25 ml of serum-free RPMI.

### Isolation of CD11b+ cells

The obtained bone marrow cells were enriched for cells expressing CD11b using anti-CD11b magnetic beads (STEM CELL Technologies, Vancouver, BC), applied according to the manufacturer’s instructions. Isolated CD11b+ cells were sent for metabolomic, lipidomic, and transcriptomic analyses.

### Lipid and metabolite extraction

Lipids and metabolites were extracted from cells using MPLEx (Metabolite, Protein, Lipid Extraction) (21, 22). First, 1 mL cold (-20 °C) chloroform:methanol working mix (prepared 2:1, v/v) was added into a chloroform-compatible 2 mL Sorenson MulTI™ SafeSeal™ microcentrifuge tube (Sorenson Bioscience, Salt Lake City, UT) inside an ice-block. The sample and cold water were then added to make a final ratio of 8:4:3 chloroform:methanol:water/sample, vortexed, allowed to incubate in the ice block for 15 mins with 2000 rpm shaking, and finally sonicated in an ice water bath sonicator. Afterwards, the samples were centrifuged at 12,000 x *g* for 10 mins to separate the polar and non-polar phases and the protein interlayer. The upper polar phase and a portion of the lower non-polar phase were allotted to metabolite samples, with the remaining lower non-polar phase used for lipid samples. The whole process was repeated once on the protein interlayer to ensure complete removal of metabolites and lipids. The protein was then washed with 500 µl of cold 100% methanol and centrifuged for five minutes to pellet the protein, and the wash layer was added to the metabolites vial. Metabolite and lipid samples were dried in a speed vac, and 500 µl 2:1 (v:v) cold chloroform:methanol was added to the lipids, which were ultimately stored along with the dry metabolites at -20 °C until analysis.

### Lipidomics analysis

Total lipid extracts (TLEs) were analyzed using reversed-phase LC-ESI-MS/MS using a Waters Aquity UPLC H class system (Waters Corp., Milford, MA) coupled with a Velos Pro Orbitrap mass spectrometer (Thermo Scientific, San Jose, CA). Prior to analysis, TLEs were evaporated and then reconstituted in 5 µl chloroform and 45 µl of methanol. For analysis, 10 µl of this reconstitution were injected onto a Waters LC column (CSH 3.0 mm x 150 mm x 1.7 µm particle size) maintained at 42 °C. Lipid species were separated using a 34 min gradient elution at a flow rate of 250 µl/min. Mobile phase A and B respectively consisted of acetonitrile/H_2_O (40:60) containing 10 mM ammonium acetate and acetonitrile/isopropanol (10:90) containing 10 mM ammonium acetate. The full gradient profile was as follows (min, %B): 0, 40; 2, 50; 3, 60; 12, 70; 15, 75; 17, 78; 19, 85; 22, 92; 25, 99; 34, 99; and 34.5, 40.

The UPLC system used a Thermo HESI source coupled to the mass spectrometer inlet. The MS inlet and HESI source were maintained at 350 °C with a spray voltage of 3.5 kV and sheath, auxiliary, and sweep gas flows of 45, 30, and 2, respectively. Each TLE was analyzed in both positive and negative ion mode in separate analyses. Lipids were fragmented by higher- energy collision dissociation (HCD) and collision-induced dissociation (CID) using a precursor scan of m/z 200-2000 at a mass resolution of 60k followed by data-dependent MS/MS of the top four most abundant precursor ions. An isolation width of 2 m/z units and a maximum charge state of 2 were used for both CID and HCD scans. Normalized collision energies for CID and HCD were 35 and 30, respectively. CID spectra were acquired in the ion trap using an activation Q value of 0.18, while HCD spectra were acquired in the Orbitrap at a mass resolution of 7.5k and a first fixed mass of m/z 90.

Lipid identifications were made using MS-DIAL v4.92 (23) for peak detection, identification and alignment. The tandem mass spectra and corresponding fragment ions, mass measurement error, and aligned chromatographic features were manually examined to remove false positives. Relative quantification was performed by calculating peak areas on extracted ion chromatograms of precursor masses. Features detected in at least three of the five replicates were retained.

### Metabolomics analysis

Dried extracts were chemically derivatized using a modified version of the protocol used to create FiehnLib (26). Briefly, dried metabolite extracts that had been stored at -80l°C were dried again to remove any residual water. To protect carbonyl groups and reduce the number of tautomeric isomers, 20 µl of methoxyamine in pyridine (30 mg ml^-1^) were added to each sample, followed by vortexing for 30 s and incubation at 37 °C with vigorous shaking (1,000 rpm) for 90 min. The sample vials were then inverted once to capture any condensation of solvent at the cap surface, followed by a brief centrifugation at 1,000 × *g* for 1 min. To derivatize hydroxyl and amine groups to trimethylsilylated (TMS) forms, 80 µl of *N*-methyl-*N*- (trimethylsilyl)trifluoroacetamide (MSTFA) with 1% trimethylchlorosilane (TMCS) were then added to each vial, followed by vortexing for 10 s and incubation at 37 °C with shaking (1,000 rpm) for 30 min. Again, the sample vials were inverted once, then centrifuged at 1,000 × *g* for 5 min. The samples were finally allowed to cool to room temperature and analyzed the same day. Analyses were performed on an Agilent 8890 GC coupled to a 5977B MSD (Agilent Technologies); samples were run in random order for each experiment. An HP-5MS column (30 ml×l0.25 mml×l0.25 µm; Agilent Technologies) was used for untargeted metabolomics analyses. The sample injection mode was splitless, and 1 µl of sample was injected. The injection port temperature was held at 250 °C throughout the analysis. The GC oven was held at 60 °C for 1 min after injection, after which the temperature was increased to 325 °C by 10 °C min^-1^, followed by a 5 min hold at 325 °C (27). The helium gas flow rate for each experiment was determined by the Agilent Retention Time Locking function based on analysis of deuterated myristic acid; all were in the range of 0.45-0.5 ml min^-1^. Data were collected over the mass range 50-550 m/z. A mixture of fatty acid methyl esters (C8-C28) was analyzed once per day together with the samples to allow for retention index alignment during subsequent data analysis.

GC-MS raw data file processing was performed using the Metabolite Detector software and metabolites were identified by matching experimental spectra and retention indices to a PNNL-augmented version of the GC-MS Metabolomics Library (26). All identifications were manually validated to reduce deconvolution errors and to eliminate false identifications. The NIST 20 GC-MS spectral library and Wiley 11th Edition GC-MS library were also used to cross- validate the spectral matching scores obtained using the Agilent library and to provide identifications of unmatched metabolites.

GC-MS raw data file processing was performed using the MS-DIAL v4.92 (23) software and metabolites were identified by matching experimental spectra and retention indices to a PNNL-augmented version of the GC-MS Metabolomics Library (26). All identifications were manually validated to reduce deconvolution errors and to eliminate false identifications.

Statistical analysis of metabolomics and lipidomics data was performed using the PMart web application (28). Data was log2-transformed and normalized via global median centering. Statistical comparisons were performed using ANOVA with a Holm test correction (29).

### Mitochondrial respiration of bone marrow cells

Oxidative phosphorylation of bone marrow cells was assessed as we described before (18). In brief, bone marrow cells were harvested from the femur and tibia of euthanized mice as previously reported (20). The day before the assay, a Seahorse sensor plate (Agilent) was calibrated overnight in a non-CO_2_ incubator. On the day of the assay, frozen bone marrow cells were thawed, counted according to staining with trypan blue, plated at 200,000 cells/well in a poly-D-lysine-coated plate, and let to rest at 37 °C for one hour in RPMI. The medium was then replaced with supplemented Seahorse basal medium and the plates were calibrated in a non- CO_2_ incubator for 45 min prior to the experiment. Drugs were added to cartridge plates at the following final concentrations: for the ATP assay, 2 mM oligomycin, 0.5 mM rotenone, and 0.5 mM antimycin A; and for the Mito Stress Assay, 2.0 mM oligomycin, 2.0 mM FCCP, 0.5 mM rotenone, and 0.5 mM antimycin A. The ATP Assay and Mito Stress Assay programs were run on a Seahorse XFe96 (Agilent, Santa Clara, CA) as we previously described (30).

### RNA isolation from CD11b+ cells

Total RNA was isolated from the purified CD11b+ cells using the RNeasy mini kit (Qiagen, Redwood City, CA), and the yield was quantified with a NanoDrop™ Spectrophotometer (Thermo Fisher Scientific, Waltham, MA). RNA integrity and size distribution were assessed using a 2100 Bioanalyzer (Agilent Technologies) and an RNA 6000 Nano kit.

### RNA library preparation, sequencing, and data pre-processing

Sequencing assays were performed in the OHSU Integrated Genomics Lab. Libraries were prepared from 200 nanograms of total RNA using the TruSeq Stranded mRNA kit (Illumina). Briefly, RNA was converted to cDNA using random hexamers. Synthesis of the second strand included the addition of dUTP, which enforced the stranded orientation of the libraries by blocking amplification off the second strand during the first round of PCR. Amplified libraries were profiled using a 4200 TapeStation (Agilent) and a D1000 Screen Tape. Quantitative PCR of the libraries was performed using an NGS Library Quantification Kit (Roche/Kapa Biosystems) and a QuantStudio 3 Real Time PCR Workstation (Thermo/ABI). The sequencing assay was run on a NovaSeq 6000 (Illumina) sequencer. Base calling was performed by RTA v3.4.4. Fastq files were assembled using bcl2fastq (Illumina), v2.20.0.422, and FastQC v0.12.1 (http://www.bioinformatics.babraham.ac.uk/projects/fastqc/) was applied to inspect the quality of the obtained sequences.

### Sequence alignment

Before aligning each fastq file to the mouse genome, reads were trimmed using Trimmomatic version 0.36 (31) with reference to the ’TruSeq3.fa’ adapter sequence file. The specific parameters applied were as follows: ILLUMINACLIP:TruSeq3.fa:2:30:10:2:keepBothReads, LEADING:3, TRAILING:3, SLIDINGWINDOW:4:15, and MINLEN:36. Subsequently, the trimmed reads were aligned to the *Mus musculus* genome assembly GRCm38 using the STAR aligner (32), version 2.5.3a, with the parameter ’outFilterMismatchNmax’ set to 2 to specify the number of mismatched bases permitted. STAR has been shown to perform well compared to other RNA-seq aligners (33). Gene counts were subsequently generated by the STAR aligner.

### Statistical analysis of RNA-sequencing data

Gene-level raw counts were filtered to remove genes with extremely low counts in many samples, following published guidelines (34); subsequently, counts were normalized using the trimmed mean of M-values method (35) and transformed to log-counts per million with associated observational precision weights using the voom (36) method. Surrogate variable analysis with primary variables of phenotype and sex was used to estimate and remove unmeasured sources of heterogeneity (37). Gene-wise linear models with said primary variables and three surrogate variables were employed for differential expression analyses using limma with empirical Bayes moderation (38) and false discovery rate (FDR) adjustment using the Benjamini and Hochberg method (39). Differential expression data were analyzed using the Ingenuity Pathway Analysis software (QIAGEN Inc., https://digitalinsights.qiagen.com/products-overview/discovery-insights-portfolio/analysis-and-visualization/qiagen-ipa/), with a stringent cutoff for significant molecules of FDR *p* < 0.2 and |log2(foldchange)| > 0.585.. The background reference set comprised all genes in the differential expression analysis.

### Data availability

The metabolomics and lipidomics data are available at the NIH Common Fund’s National Metabolomics Data Repository (NMDR) website, the Metabolomics Workbench, https://www.metabolomicsworkbench.org, where they have been assigned Study ID ST002803. RNA-sequencing data were submitted to Gene Expression Omnibus, the accession number is GSE275254.

## RESULTS

### Mouse model of maternal obesity

Six-week-old females were placed either on a regular (13% kcal from fat) or a high-fat (45% kcal from fat) diet (RD and HFD, respectively) (**Fig. 1A**). After eight weeks of dietary intervention, the HFD-fed females were bred to RD-fed males. As we have previously reported, no differences in body weight were observed between RD- and HFD-fed mothers-to-be prior to breeding (19, 40). However, visceral adiposity was increased in HFD-fed mothers when compared with RD-fed mothers (18, 19)

**Figure 1.**
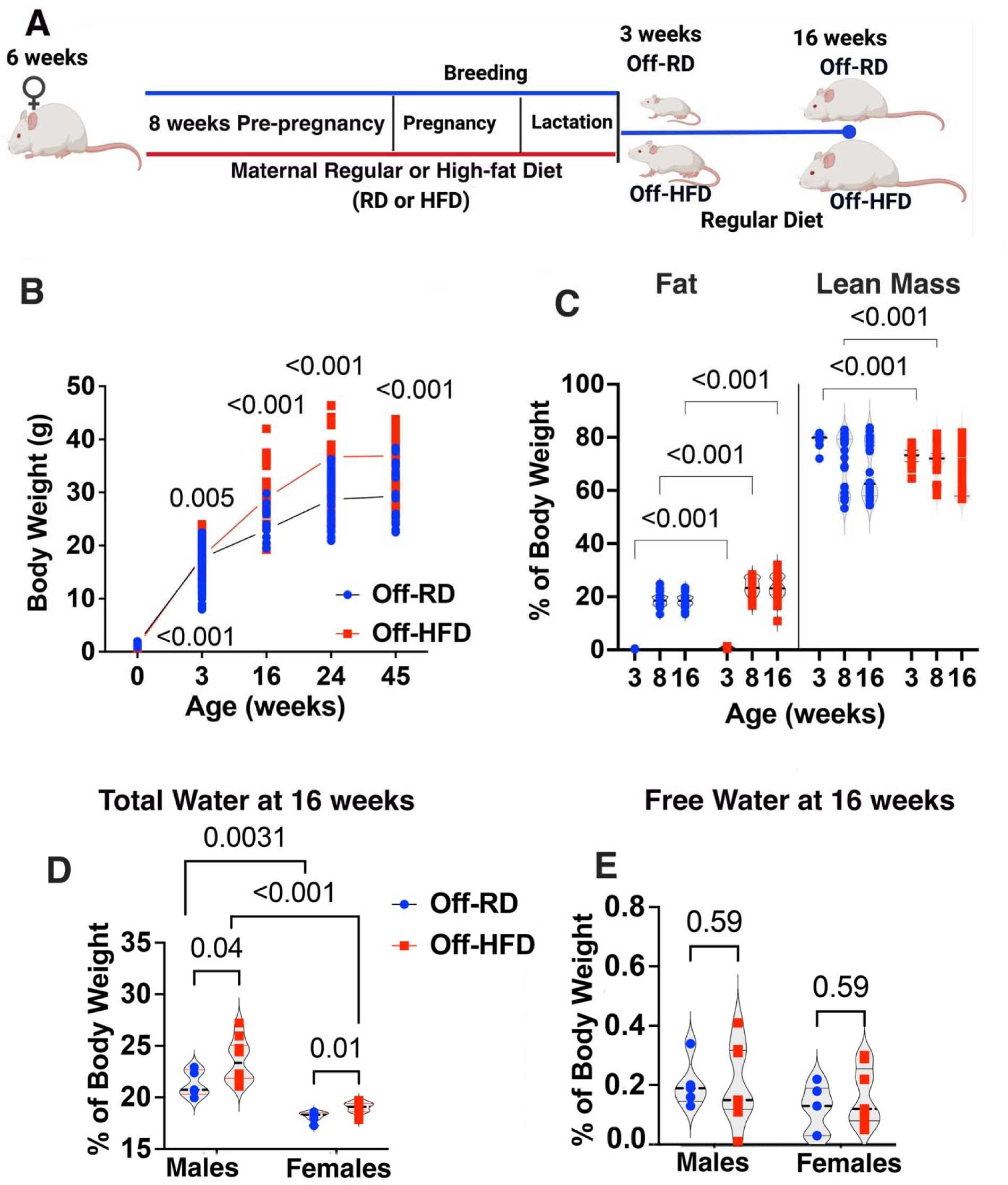
Body weight gain and body composition in Off-RD and Off-HFD. **A**, Mouse model of maternal obesity. Illustration created with BioRender.com. **B**, Dynamics of body weight gain in Off-RD and Off-HFD mice from weaning (3 weeks) through 6, 16, 24, and 45 weeks of age. Data were analyzed by two-way ANOVA followed by multiple unpaired *t*-tests. N=20- 44/group of maternal diet/age, *p*-values are shown for Off-HFD vs. Off-RD at each time point. Solid lines indicate average body weight (blue, Off-RD; red, Off-HFD). Data were analyzed by two-way ANOVA followed by multiple unpaired *t*-tests with FDR<0.05. **C**, Body fat and lean mass of 3-, 8-, and 16-week-old Off-RD and Off-HFD mice, expressed as a percentage of body weight. N=12-24/group of maternal diet/sex. Data from males and females were pooled. **D-E**, Fetal-sex-specific changes in levels of **D**, total water and **E**, free water in 16-week-old Off-RD and Off-HFD mice, measured by echoMRI and expressed as a percentage of body weight. Data were analyzed by two-way ANOVA followed by multiple unpaired *t*-tests. N=6-9/sex/group of maternal diet. *P*-values are shown on the graphs.

Consistent with our previous report (19), the offspring of HFD-fed mothers (Off-HFD) were born with lower birth weights, but caught up during lactation, and from weaning (three weeks) and up to 45 weeks of age were heavier compared with the offspring of RD-fed mothers (Off-RD, *p*<0.05, **Fig. 1B**). Despite being fed RD only, offspring of HFD-fed mothers continued to gain significant amounts of fat mass throughout their lifetime. Echo MRI analysis of offspring adiposity, lean mass, and water content showed significantly increased body fat percentage in the Off-HFD at 3, 8, and 16 weeks of age compared with age- and sex-matched Off-RD mice (**Fig. 1C**, *p*<0.05). At the same time, lean mass, a measure of muscle tissue in all body parts containing water, was decreased in three- and eight-week-old Off-HFD vs. Off-RD. Interestingly, 16w old animals showed a decrease in lean mass when compared to the 8w old animals in both Off-HFD and Off-RD. Despite the significant increase in adiposity, the percentage of lean mass remained unchanged in 16-week-old Off-HFD vs. Off-RD probably because of the decrease in lean mass from 8w to 16w. We then compared total water, which includes both free water and the water contained in lean mass, between the groups. The percentage of total water was significantly increased in 16-week-old male and female Off-HFD vs. Off-RD (**Fig. 1D**), while free water remained unchanged (**Fig. 1E**), suggesting an accumulation of water in internal organs. These results indicate that lipid content of the body remains high from the very early age in Off- HFD when compared to the Off-RD.

### Changes in bone marrow cell lipidome and energy metabolism

We have previously reported significant metabolic changes in the bone marrow of offspring of HFD-fed mothers (18). Since we observed an increase in fat content of the body, we hypothesized that lipid metabolism is also perturbed in the bone marrow cells in Off-HFD vs Off- RD. Thus, we explored pathways through which maternal obesity affects bone marrow lipid metabolism. We performed lipidomics profiling of bone marrow cells from three-week-old mice (**Fig. 2**). As our previous study established that exposure to maternal obesity is associated with critical changes in myeloid cell population, we also analyzed CD11b^+^ cells from the bone marrow of these animals using metabolomics, lipidomics, and transcriptomic approaches (**Fig. 2**).

**Figure 2.**
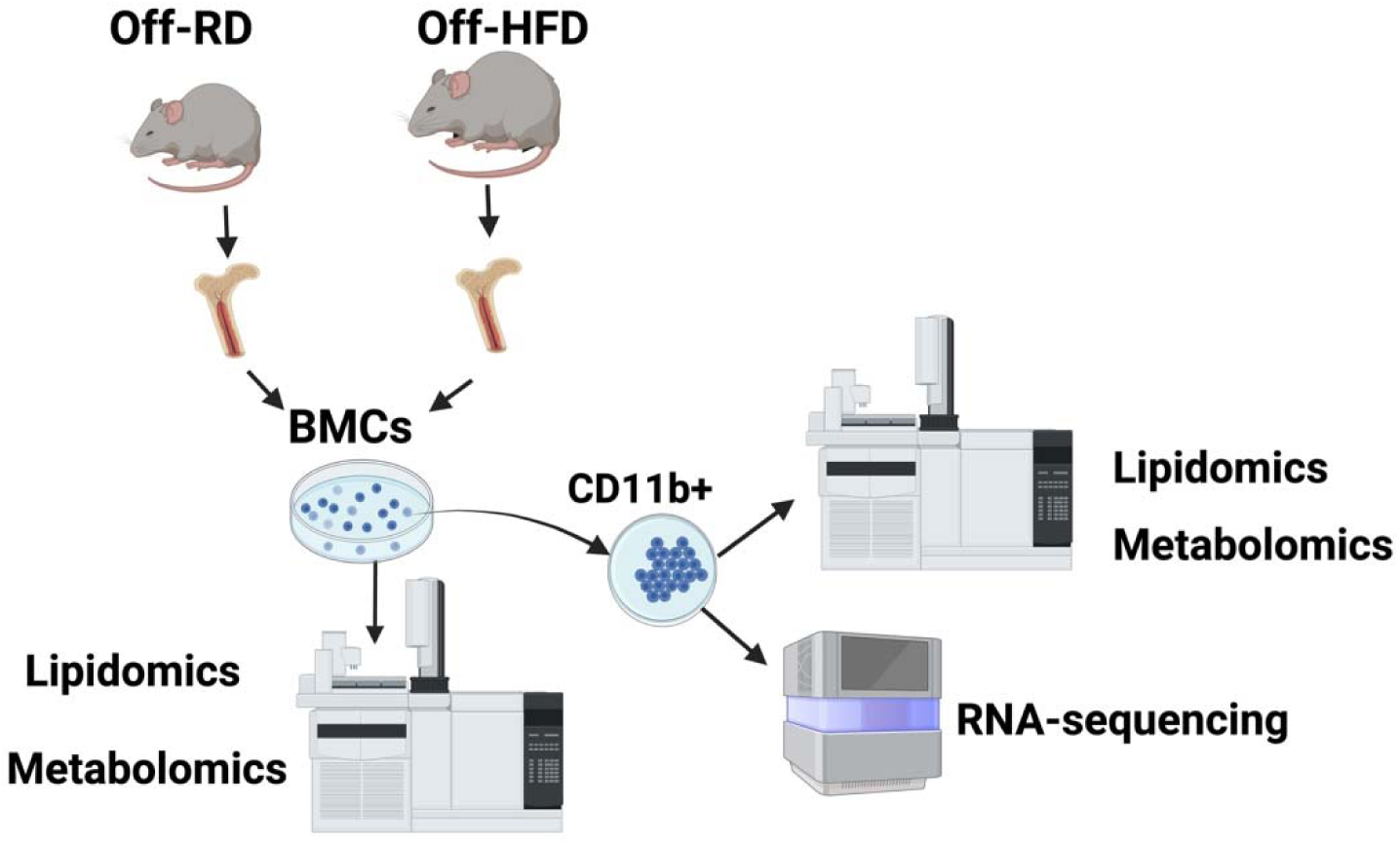
Schematic of the experimental approach: isolation of bone marrow cells (BMCs) and purification of CD11b+ cells from the offspring of mothers fed a regular (Off- RD) or a high-fat diet (Off-HFD). Bone marrow cells were isolated from the femurs of three- weeks-old mice and sent for metabolomics and lipidomic analyses. In addition, CD11b+ cells were purified from bone marrow cells and submitted for metabolomics, lipidomics and RNA- sequencing analyses. Illustration created with BioRender.com.

Bone marrow cell lipidomics revealed broad, dynamic changes in lipid profiles as result of exposure to maternal obesity (**Fig. 3**). Overall, 113 differentially expressed lipids were identified in male and female Off-HFD vs. Off-RD (false discovery rate [FDR] calculated by the Benjamini and Hochberg method <0.05, *p*<0.05). These were divided into four classes according to their lipid backbones (**Fig. 3A**): diacylglycerols (DAGs), triacylglycerols (TAGs), sphingomyelins, and phospholipids. Notably, DAGs and TAGs displayed the most significant differential regulation, with a predominant increase and decrease in abundance respectively in Off-HFD vs. Off-RD mice (**Figs. 3B and 3D**). Among phospholipids, lyso-phosphatidylcholines (LPCs) showed a significant decrease in Off-HFD mice (*p*<0.001), whereas phosphatidylcholines (PCs) and phosphatidylethanolamines (PEs) were increased (*p*<0.001 for both, **Fig. 3C**). Notably, we observed reduced abundance of several PC and PE plasmalogens, a subclass of glycerophospholipids for which reduced levels have been observed in neurodegenerative (41), cardiovascular (42), and chronic inflammatory diseases (43), and also in aging (44). The affected plasmalogens included PC P-38:5, PC P-38:4, PC P-40:7, PC P- 38:3, PE P-38:4, and PE P-40:5. Sphingolipid remodeling was evident in reduced levels of sphingomyelins in the bone marrow of Off-HFD vs. Off-RD mice (**Fig. 3E**).

**Figure 3.**
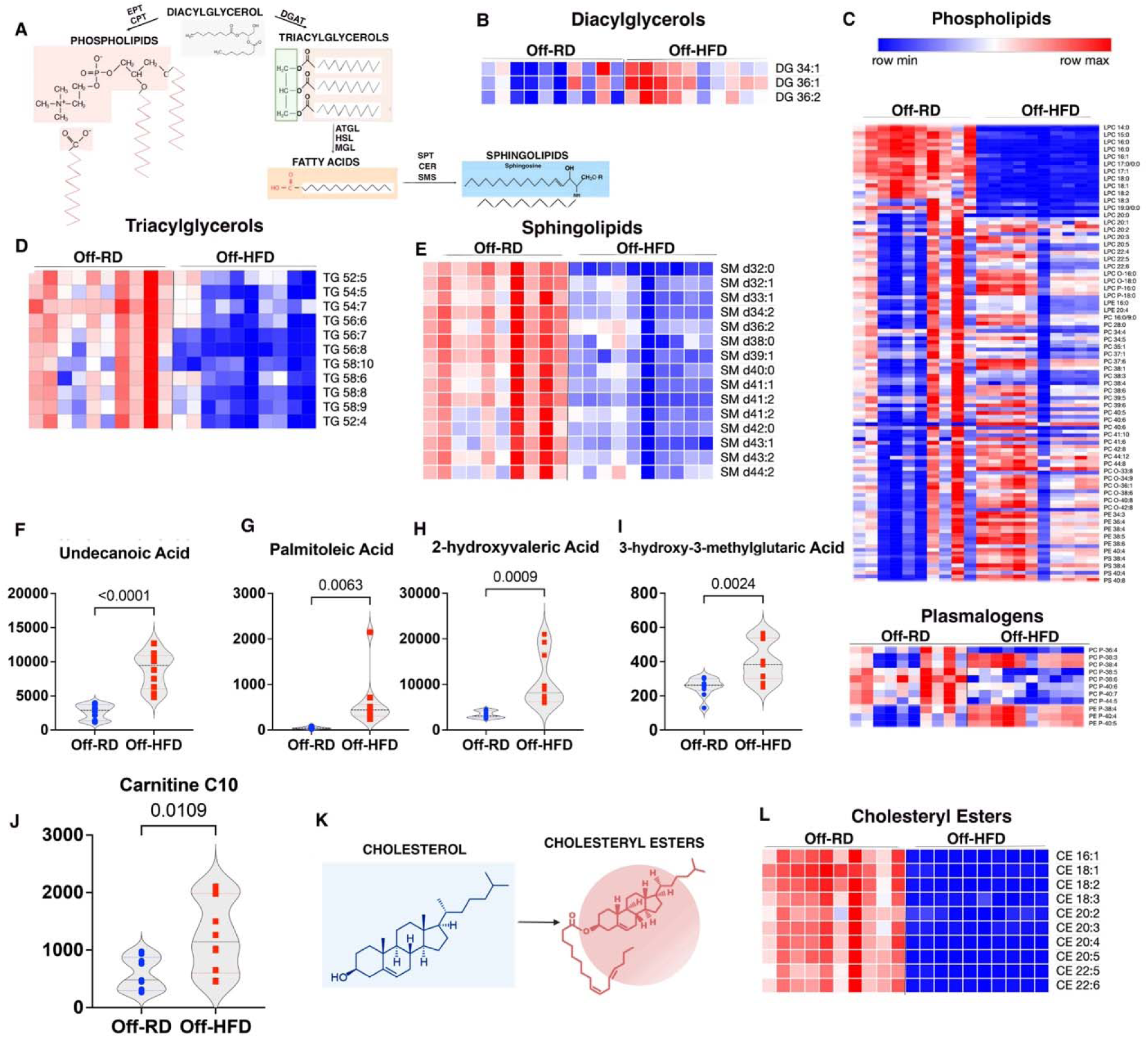
Dysregulations in bone marrow lipid species in newly-weaned Off-HFD mice. **A**, General schematic of metabolic pathways for the synthesis of phospholipids, triacylglycerols, and sphingolipids from diacylglycerols. **B-E**, Heat maps showing average levels of particular lipid species in bone marrow cells. More intense red tones represent more positive values, while blue tones represent negative values. Lipids shown are: **B**, diacylglycerols (DG); **C**, phospholipids: lysophosphatidylcholines (LPC), phosphatidylcholines (PC) including plasmalogens, phosphatidylethanolamines (PE), and phosphatidylserines (PS); **D**, triacylglycerols (TG); and **E**, sphingomyelins (SM). **F-I**, Changes in saturated, unsaturated and hydroxy fatty acids: **F**, undecanoid; **G**, palmitoleic; **H**, 2-hydroxyvaleric; and **I**, 3- hydroxymethylglutaric. **J**, Change in C10 carnitine. **K**, Schematic of the conversion of free cholesterol to cholesteryl esters. **L**, Heat map showing average levels of different species of cholesteryl esters in bone marrow cells. N-10/group of maternal diet/sex, 40 animals in total. *p*-values are either shown or *p*<0.05, Off-HFD vs. Off-RD. EPT, Ethanolaminephosphotransferase; CPT, CPT1, cholinephosphotransferase; DGAT, diglyceride acyltransferase; ATGL, adipose triglyceride lipase; HSL, hormone-sensitive lipase; MGL, monoacylglycerol lipase; SPT, serine palmitoyltransferase complex; CERs, ceramide synthases; SMS, sphingomyelin synthases. Illustrations created with BioRender.com.

Decreased TAGs could be explained by enhanced oxidative phosphorylation, that we have reported before (18). Consistent with this hypothesis, we found four fatty acids significantly increased by maternal HFD: undecanoid acid, a saturated fatty acid containing eleven carbons (**Fig. 3F**); palmitoleic acid (**Fig. 3G**), an unsaturated fatty acid previously reported to be increased in obese children and adults (45, 46); and two hydroxy fatty acids, 2-hydroxyvaleric acid and 3-hydroxy-3-methylglutaric acid (**Fig. 3 H-I**). Transfer of long-chain fatty acids into the mitochondrion is a rate-limiting step in fatty acid oxidation, mediated by carnitines. As expected, our data revealed significantly increased C10 carnitine in bone marrow cells from Off-HFD vs. Off-RD mice (**Fig. 3J**).

Cholesterol is critical for immune cell proliferation and intracellular metabolism (47), being responsible for mitochondrial membrane fluidity, permeability, and hence respiration (48). It can exist in two forms: the non-esterified or free form and cholesteryl ester (CE). Free cholesterol is biologically active and can have harmful effects on cells (49), while CE is a protective form that can be transported in lipoprotein particles to tissues that use cholesterol or stored in the liver in lipid droplets (**Fig. 3K**) (50). Insufficient conversion of cholesterol to CEs has been previously described in the blood of patients with cardiovascular diseases (51). Our data revealed a significant decrease in CE abundance in the bone marrow of Off-HFD compared with Off-RD mice (**Fig. 3L**). Furthermore, upon performing pathway enrichment analysis using REACTOME, KEGG, and the MetaboAnalyst software (**Table 1**), the differential metabolites were mapped onto 15 pathways related to cholesterol and lipid metabolism. These comprised: HDL, LDL, VLDL, and chylomicron remodeling, assembly, and clearance; vesicle- mediated transport, lipid particle organization, membrane trafficking, chylomicron clearance and assembly, the immune system, glycerophospholipid biosynthesis; and phospholipid metabolism.

**Table 1.**
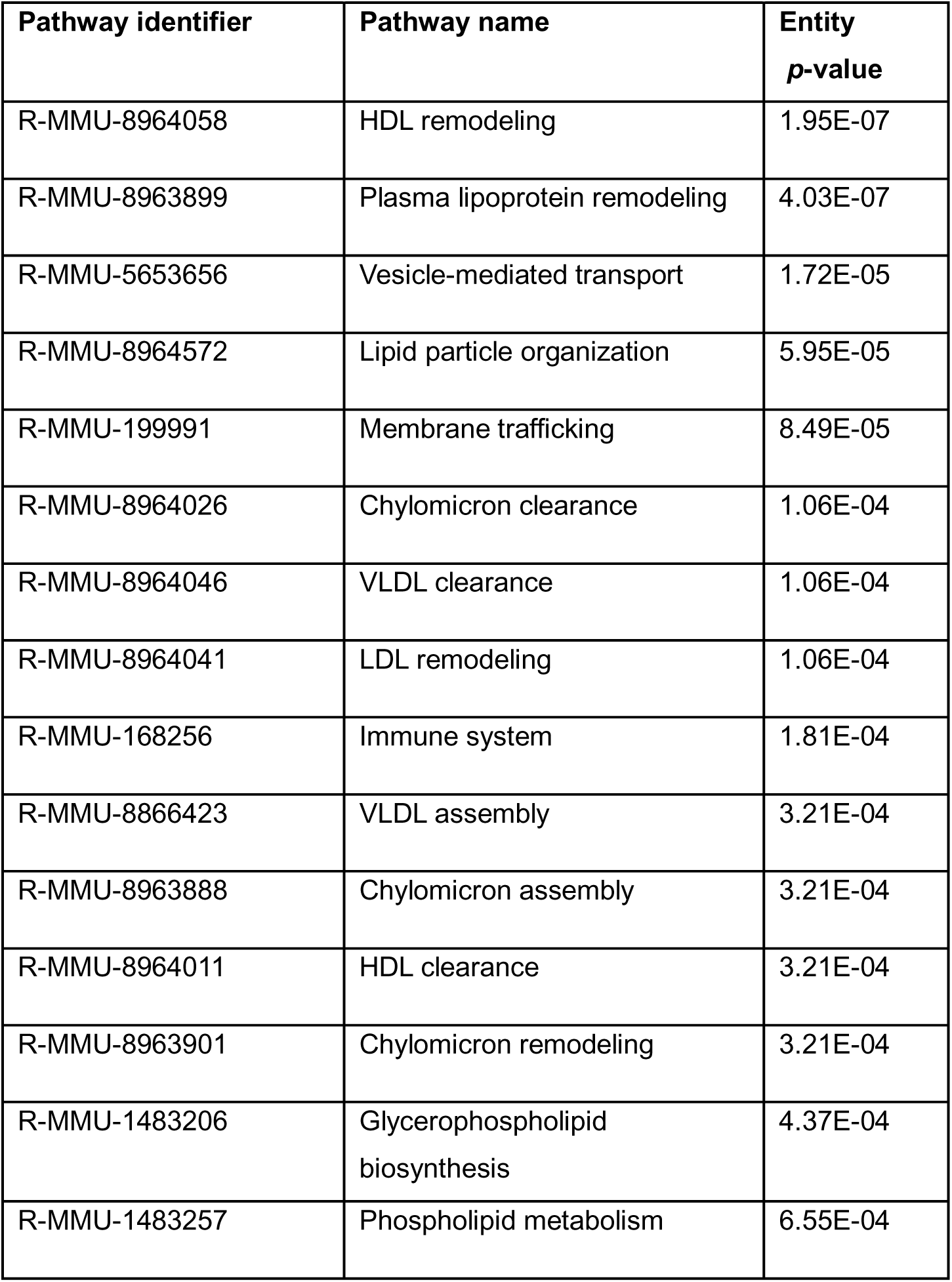
Lipidomics and metabolomics enrichment analysis results for canonical pathways in REACTOME, KEGG, and MetaboAnalyst in bone marrow cells of Off-HFD vs. Off-RD mice.

Prolonged accumulation of palmitoleic acid has previously been linked to increased oxygen consumption (52). At the same time, published evidence suggests that abnormally low levels of cholesterol and accumulation of 3-hydroxy-3-methylglutaric acid are associated with progression of mitochondrial toxicity and dysfunction in both patients and animal models (53, 54). Considering these long-term effects, we next hypothesized that oxidative phosphorylation is decreased in adult Off-HFD mice. To address this hypothesis, we used a Seahorse Analyzer to measure mitochondrial respiration in bone marrow cells isolated from Off-RD and Off-HFD mice at 16 weeks (**Fig. 4A**). We have previously reported significant metabolic and respiratory abnormalities in Off-HFD mice at this age (19, 55). In contrast to Off-RD mice, which displayed no age-dependent changes, the oxidative phosphorylation in Off-HFD was increased at three weeks of age (18), but significantly diminished at 16 weeks of age, which was evidenced by reduced production of ATP (**Fig. 4B**), basal respiration (**Fig. 4C**), ATP-induced and maximal respiration (**Fig. 4D**), and spare capacity (**Fig. 4E**), the difference between maximal and basal mitochondrial oxygen consumption rates.

**Figure 4.**
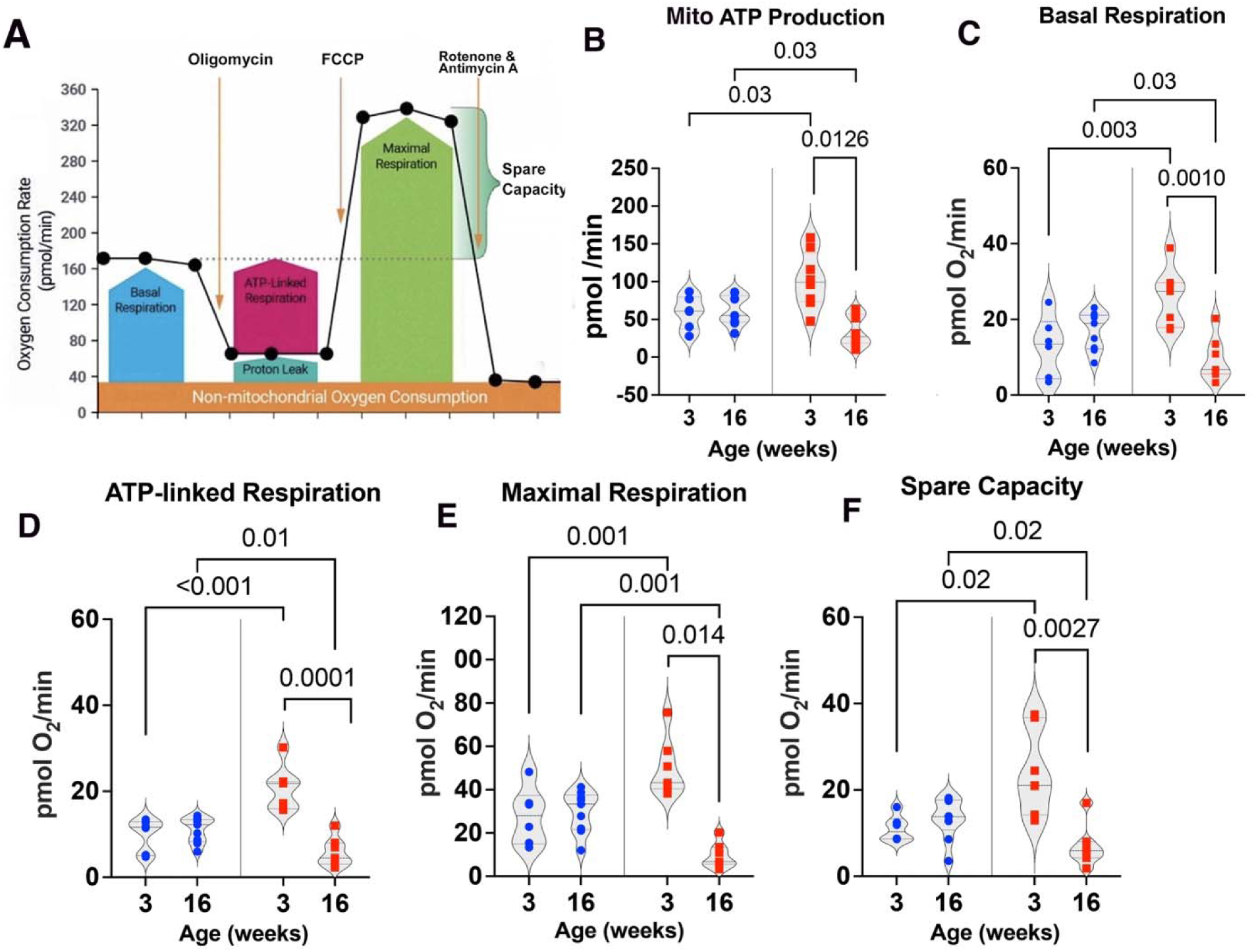
Age-dependent differences in mitochondrial respiration in bone marrow cells from Off-HFD and Off-RD mice. Oxygen consumption rates were measured using the Seahorse XF Cell Mito Stress Test in cultured bone marrow cells from Off-RD and Off-HFD. Data from three-week-old offspring are shown for comparison only. Measurements were taken before and after treatment with oligomycin (1.0lµmol/L), FCCP (1.1lµmol/L), and antimycin A/rotenone (0.5 µmol/L each), and the data were analyzed with the Wave 2.6 software. **A**, Overall workflow. **B**, Rate of ATP production from oxidative phosphorylation. **C**, Basal respiration. **D**, ATP-linked respiration. **E,** Maximal respiration. **F**, Spare respiratory capacity. Data are represented as individual values with lines at mean with SEM; *p*-values are shown. Data were analyzed by three-way ANOVA, with results from males and females pooled; N=7- 9/group of maternal diet.

### Dysregulation of bone marrow myeloid cell metabolism in the offspring of obese mothers

Accumulating evidence indicates reprogramming of macrophages and their bone marrow precursors to occur in the settings of metabolic diseases (47). We have recently reported significant changes in the immune complexity of bone marrow cells, specifically CD11b+ cells, in three-week-old offspring of HFD-fed mothers (18). We therefore purified CD11b+ cells from the bone marrow of three-week-old Off-RD and Off-HFD mice using immunomagnetic positive selection and conducted metabolomic, lipidomic, and transcriptomic analyses to investigate the effect of maternal obesity on myeloid cell metabolism and gene expression. This revealed amino acid metabolism to be the most altered in myeloid cells: especially, our data revealed a significant increase in aspartic acid (**Fig. 5A-B**), and a decrease in its conversion product L- homoserine (*p*<0.05, **Fig. 5C**). It is noteworthy here that aspartic acid is a key anabolic amino acid for its proteogenic and nucleotide biosynthesis role in cells and is found to be essential for cell proliferation (56). In fact, aspartate biosynthesis has been shown to be a key function of electron transport chain in proliferating cells (57). Other amino acids and metabolites were changed only in a sex-specific manner. For example, amino acids L-threonine, L-valine, and pyroglutamic acid showed significant increases in female vs. male offspring of both maternal diet groups (**Supplemental Figure 1A-D**). In contrast, Krebs cycle metabolites L-malic and fumaric acid were increased in female Off-RD but showed no significant sexual dimorphism in Off-HFD mice (**Supplemental Figure 1A-D**).

**Figure 5.**
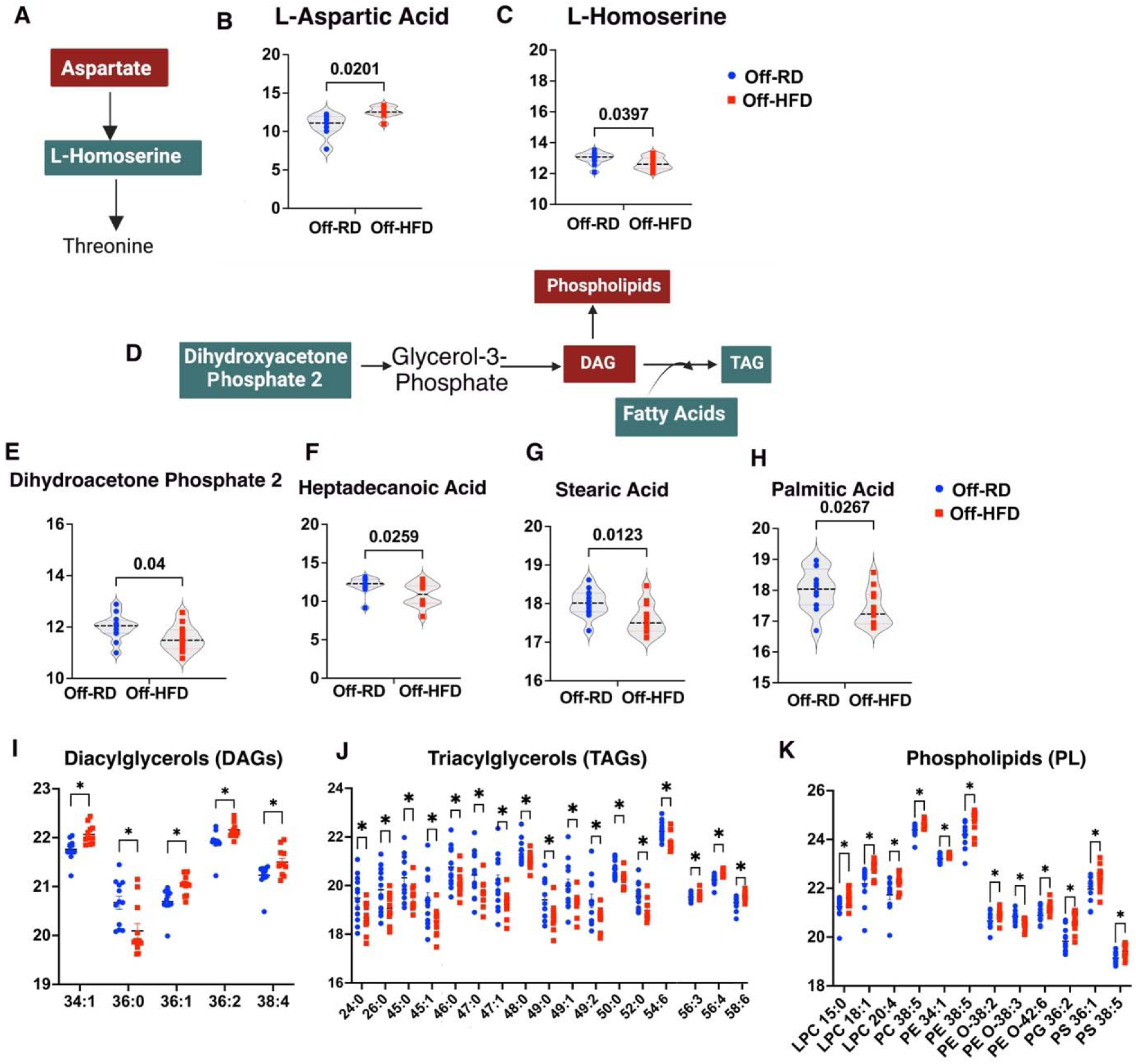
Metabolomic and lipidomic differences in bone marrow CD11b+ cells of Off-HFD vs. Off-RD mice. **A-C**, Alteration in amino acid levels. **A,** Biosynthesis of threonine from aspartic acid and levels of the amino acids **B**, L-aspartate and **C**, L-homoserine. **D-K,** Alteration in lipid metabolism in response to maternal obesity. **D,** Schematic of triglyceride synthesis from dihydroacetone phosphate 2 to triglycerides. Significant changes were observed in levels of **E,** dihydroacetone phosphate 2 and several saturated fatty acids: **F**, heptadecanoic; **G**, stearic; and **H**, palmitic. Also altered in the myeloid cells of 3-week-old mice were: **I**, diacylglycerols; **J**, triacylglycerols; and **K**, phospholipids. Data from males and females were pooled. Data are represented as individual values. N=6/sex/group of maternal diet, *p*-values are shown in panels (A-H) and otherwise indicated as *, *p*<0.05. Illustrations in A and D created with BioRender.com.

Our data also revealed significant reduction of dihydroxyacetone phosphate, an intermediate metabolite in TAG synthesis, in the myeloid cells of Off-HFD vs. Off-RD mice (**Fig. 5D-E**), while its downstream product glucose-3 phosphate was affected in a sex-dependent manner (**Supplemental Figure 1G-H**), showing an increase in female vs. male Off-RD but not Off-HFD mice. Surprisingly, the levels of three saturated fatty acids, stearic, heptadecanoic, and palmitic acid, were significantly decreased in myeloid cells of Off-HFD vs. Off-RD (*p*<0.05, **Fig. 4F-H**). Myristic acid, another long-chain saturated fatty acid, was altered in a sex-dependent manner, having significantly lower expression in females compared with males within both maternal diet groups (**Supplemental Figure 1I**).

Consistent with bone marrow cells, lipidomics analysis of myeloid cells revealed Off- HFD to have significantly higher levels of DAGs when compared with Off-RD mice (**Fig. 4I**).

Also similar to bone marrow cells, TAG levels in CD11b+ cells were reduced in the Off-HFD group, except for three high-carbon-number TAGs that showed significant accumulation, including 56:3, 56:4, and 58:6 (**Fig. 4J**). All differentially expressed phospholipids, including LPCs, PCs, and PEs, were increased in bone marrow myeloid cells of Off-HFD vs. Off-RD mice (*p*<0.05, **Fig. 5K**).

We next compared the differentially expressed lipids shared in common between bone marrow cells (BMCs) and CD11b+ myeloid cells (**Fig. 6A**). There were five common lipids: two DAGs, 34:1 and 36:2; two TAGs, 52:0 and 58:6; and PE 36:4 (**Fig. 6B).** Overall, most lipids showed a consistent direction of change in BMCs and CD11b+ cells (**Fig. 6C-D**): affected lipids included TAGs, DAGs, glycerophosphoserines, PEs, and ceramides, suggesting that myeloid cells might drive the signal observed in BMCs for these classes. The only exception was TAG 58:6, which was decreased in BMCs and increased in myeloid cells.

**Figure 6.**
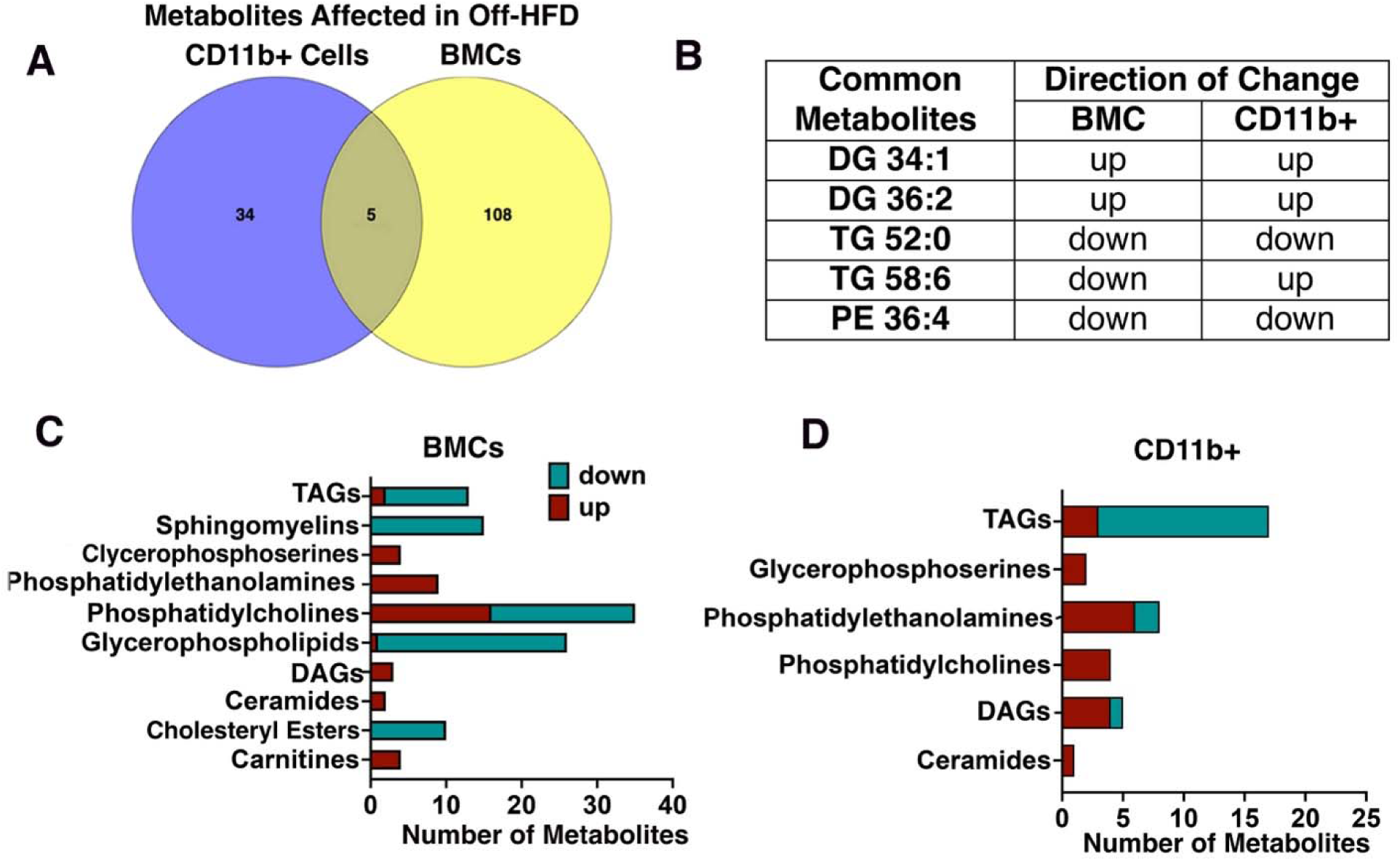
Comparison of differentially abundant lipids in bone marrow and CD11b+ cells of Off-HFD mice. **A**, Venn diagram showing numbers of unique and shared metabolites. **B**, Affected metabolites common to bone marrow and CD11b+ cells. **C-D**, Number of metabolites in each category and direction of change: **C**, in bone marrow and **D**, in CD11b+ cells. Illustrations in A, C, and D created with BioRender.com.

### Transcriptomic profiling of bone marrow myeloid cells

To link the metabolomic and lipidomic changes to gene expression, we conducted RNA- sequencing analysis on CD11b+ cell populations isolated from the bone marrow of three-week- old Off-RD and Off-HFD mice. Overall, 21 differentially expressed genes were detected that showed Benjamini and Hochberg-corrected false discovery rates (FDR) <0.2, *p*<0.05 (**Fig. 7A**), which included nine downregulated and 12 upregulated genes. We found regulators of lipid metabolism (*Lpl*), *protein processing* (*Padi4, Stfa2* and *Stfa3,Cstdc4*, *Cstdc5*, *Cstdc5*) in addition to the genes involved in immune cell trafficking and infiltration (*Jchain*, *Ighg2b*, *Padi4*, *Trio*, *Igf2bp3*, *Acvrl1*, *Dock9*, *Mmp25*), B-cell receptor signaling (*Ghg2b*, *Igkv15-103*, *Igkv16- 104*), connective tissue development and function (*CD5L*), cell-to-cell signaling (*Nrn*), amyloid beta production (*Ldlrad3*), TGFβ signaling (*Zfyve9*), (**Fig. 7B**). The upregulation of *Lpl* in myeloid cells further indicates that the utilization of fatty acids and other lipids for cell metabolism is increased in Off-HFD.

**Figure 7.**
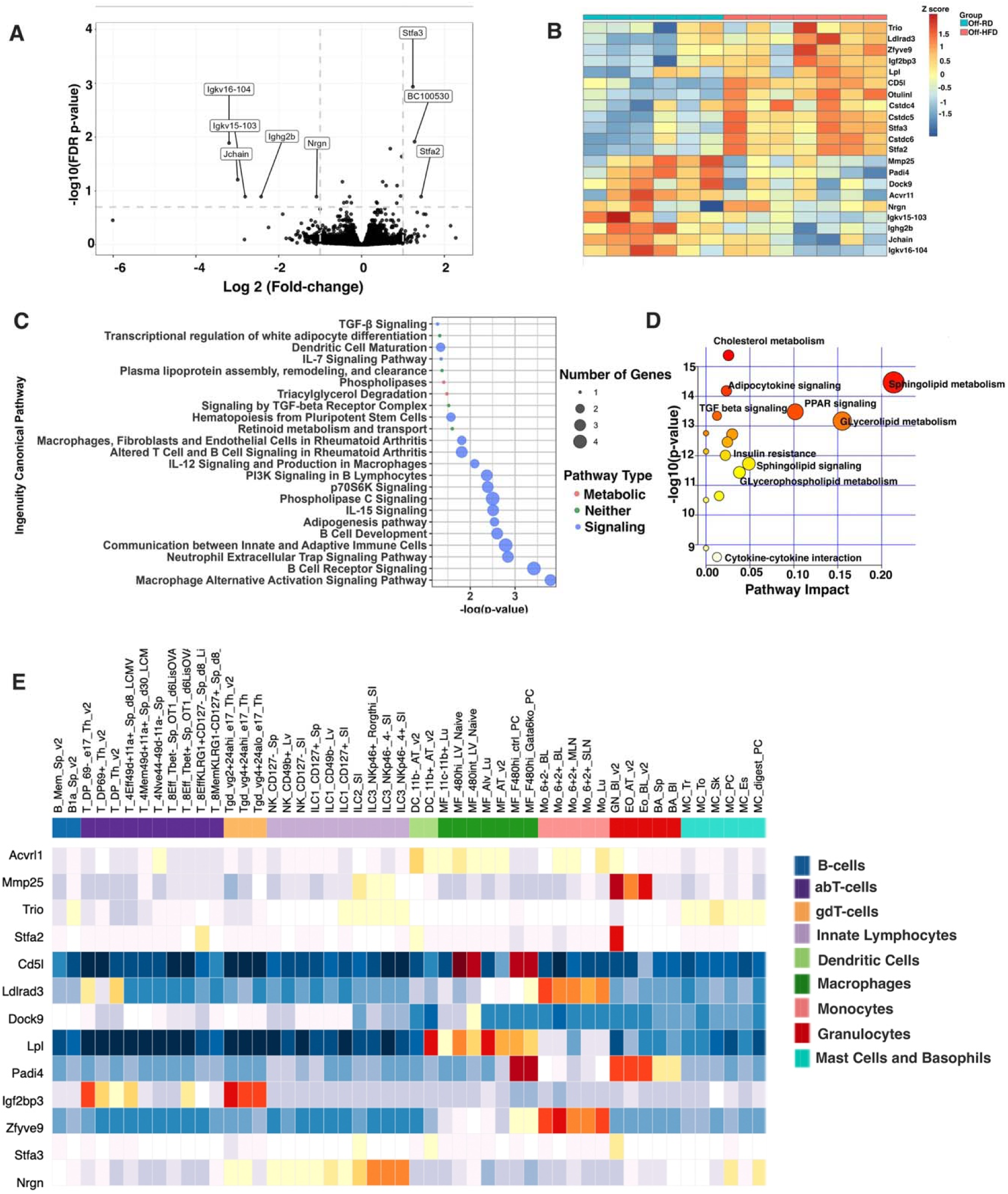
Transcriptional profiling using RNA sequencing and Ingenuity Pathway Analysis of CD11b+ myeloid cells isolated from the bone marrow of 3-week-old Off-RD and Off-HFD mice. **A**, Volcano plot showing statistical significance (FDR-corrected *p*-value) vs. magnitude of change (fold change). Dotted lines indicate the threshold for significance, FDR *p* < 0.2 and fold-change > 2. Data from males and females were pooled. **B**, Heat map for genes differentially expressed in Off-HFD vs. Off-RD. Color bar along top indicates group; N=6- 7/group. **C**, Bubble chart showing canonical pathways identified by Ingenuity Pathway Analysis as enriched in CD11b+ cells. **D**, Integration of CD11b+ myeloid cell metabolomics, lipidomics and RNA-seq data using MetaboAnalyst. **E**, Cell-type-specific expression of differentially expressed genes in ImmGen Consortium expression data. Heatmap colors correspond to expression intensity (blue, no or low expression; red, high expression). Color bar along top indicates cell type.

Next, we performed Ingenuity Pathway Analysis (IPA) to determine what cellular pathways were enriched among these differentially regulated genes. This yielded 16 causal networks (**Table 2**). Surprisingly, five of the inhibited networks (z-score<-2.0, *p*<0.05) were protein complexes involving Nuclear Receptor Subfamily 1 Group D Member 1 (NR1D1) or Rev- ErbAalpha associated with regulation of the circadian clock (NR1D1:heme:Corepressors:NPAS2 and NR1D1:heme:Corepressors: ARNTL), (NR1D1:heme:Corepressors:CLOCK) (58, 59) and epigenetic changes (NCoR/SMRT) (60). Meanwhile, activated networks (z-score >2.0, *p*<0.05) included the serpin family (SERPINF) with anti-angiogenesis function (61); the CITED2:PITX2 complex that inhibits pro-inflammatory gene expression in myeloid cells (62); the C/EBP group that regulates normal and malignant myelopoiesis (63); apelin receptor (APLNR), a member of the G protein-coupled receptor gene family that promotes hematopoiesis (64); and CDC42, which regulates the balance between myelopoiesis and erythropoiesis (65).

**Table 2.**
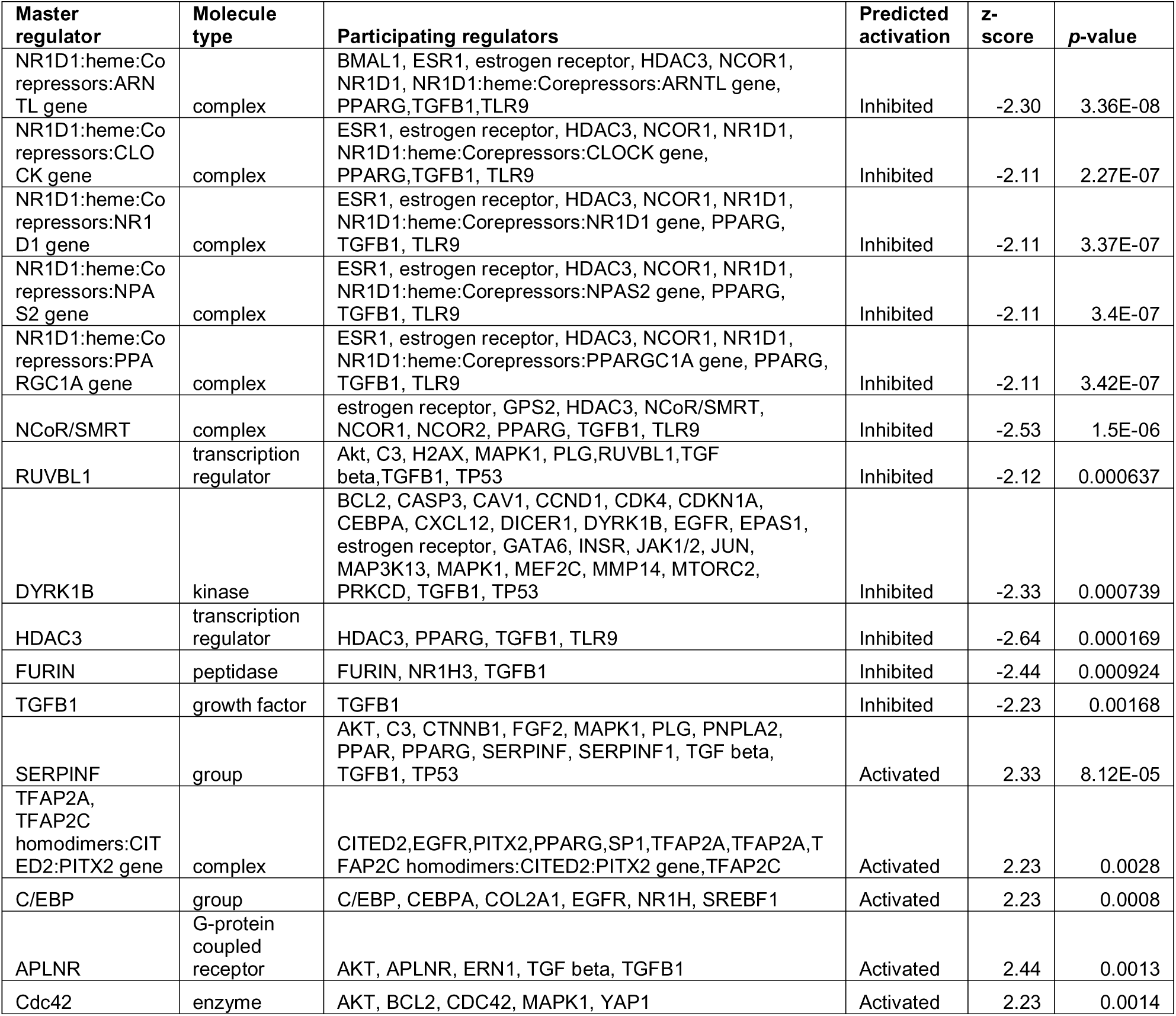
IPA enrichment results for causal networks in CD11b+ cells isolated from three-week- old Off-RD and Off-HFD mice. A network is considered activated when z-score >2.0 and inhibited when z-score <-2.0. -log10 *p*-values are shown, with -log10(*p*) > 1.3 corresponding to a *p*-value of < 0.05.

IPA also revealed several affected canonical pathways associated with differentially expressed genes (**Fig. 7C**), including macrophage alternative activation, B-cell receptors, lipid metabolism and triacylglycerol degradation, adipogenesis, communication between innate and adaptive immune cells, and TGFβ signaling The IPA-generated list of upstream regulators included four mature microRNAs (miRs) (**Table 3**): miR-33, involved in lipid metabolism and immune response during atherosclerosis (66); miR-223, regulated by lipids in myeloid cells (67); and miRs-328-3p and -3173-5p, both involved in carcinogenesis (68, 69). In addition, IPA identified six cytokines (CCL2, CX3CL1, IL17F, IL2, IL4, and IL6) as potentially dysregulated in response to maternal obesity.

**Table 3.**
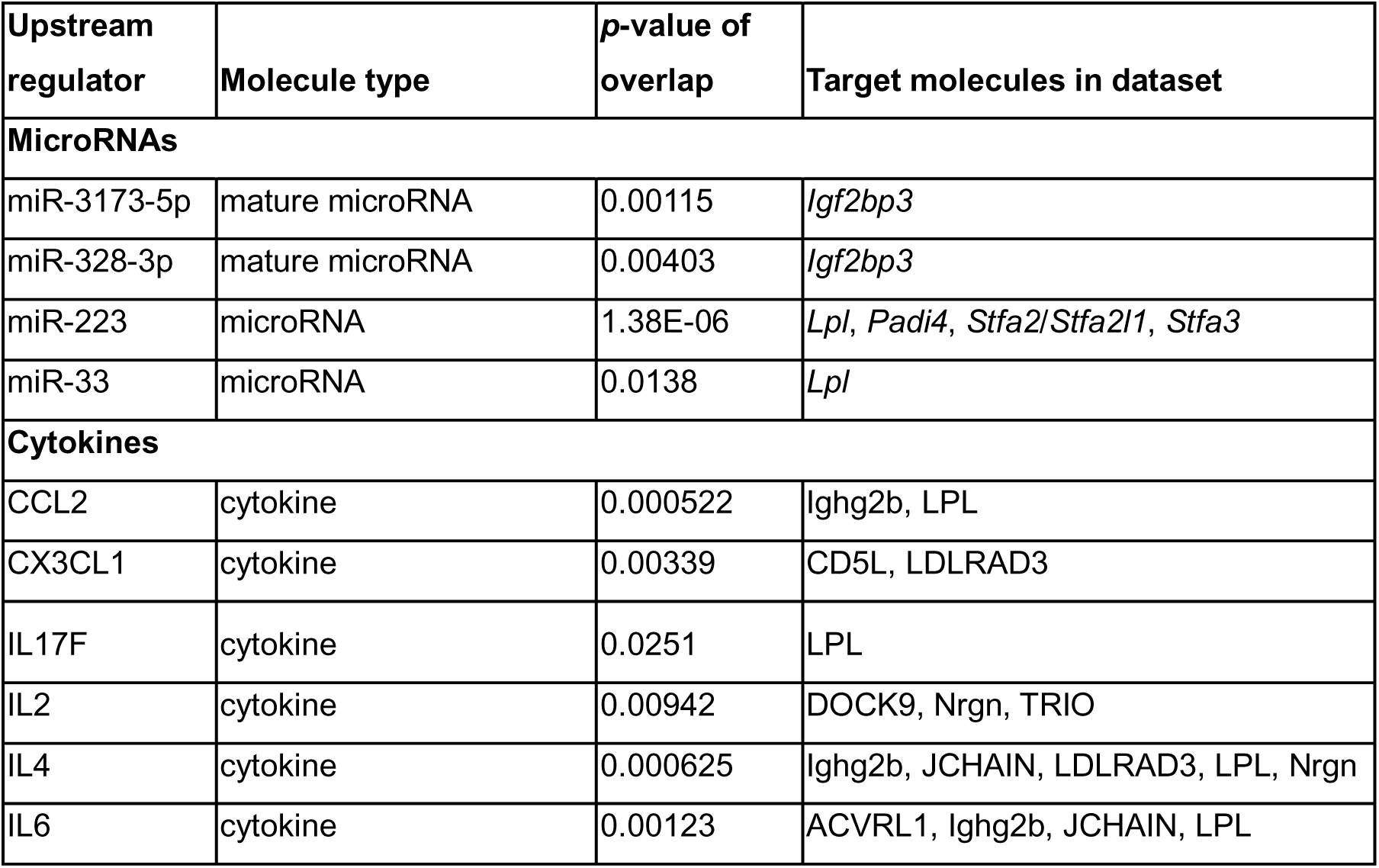
Upstream growth factors and miRNAs identified by IPA-based comparison analysis as potentially regulating genes altered in CD11b+ cells of Off-HFD mice.

Affected upstream transcription factors (**Table 4**) included YAP1, associated with immunosuppression (70); HOXA10 and CREB1 factors, regulators of normal and malignant hematopoiesis (71, 72); SP1, which has been shown *in vivo* to bind the CD11b promoter specifically in myeloid cells and is a major player in myeloid-cell-specific promoter activity; STAT6, which increases M2 polarization (73); GATA2 and GATA6, which regulate macrophage and dendritic cell function and macrophage self-renewal (74, 75); and CEBPA, a myeloid lineage enhancer (76).

**Table 4.**
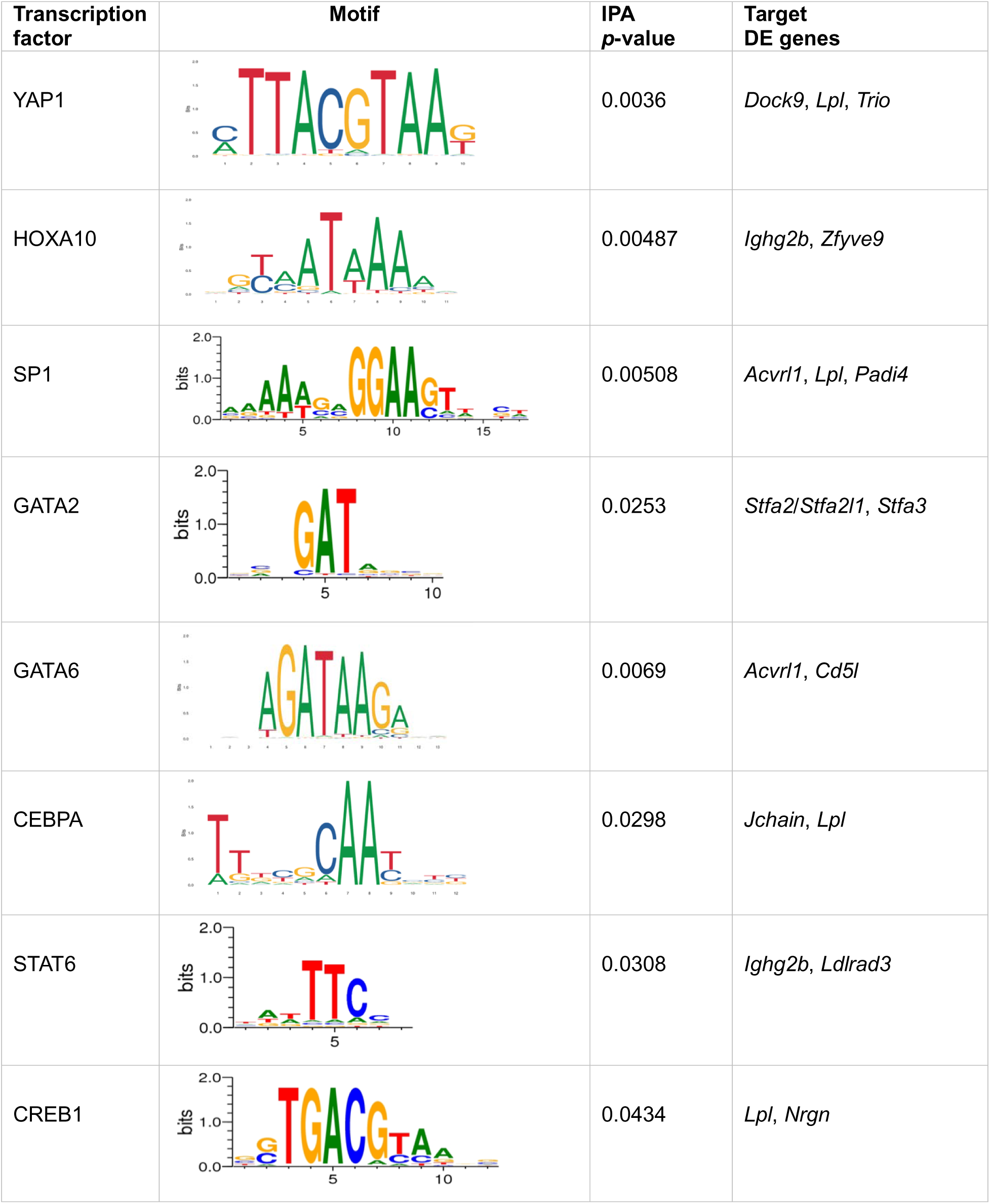
Upstream transcriptional regulators with binding motifs identified by IPA as enriched in the promoters of genes differentially expressed in CD11b+ cells of Off-HFD mice. Motif sequence logos were visualized using MotifMap.

Using MetaboAnalyst, we next integrated our lipidomics, metabolomics, and RNA- sequencing data. As shown in **Fig. 7D**, this analysis revealed several highly impacted metabolic and signaling pathways, including cholesterol and sphingolipid metabolism, insulin resistance, and the adipocytokine, PPAR, and TGFβ signaling pathways.

Finally, we set out to determine if the genes that showed differential expression were specific to myeloid cells or if they were also expressed in other immune cell populations. Towards this end, we analyzed gene expression data from sorted bone marrow cells obtained from the Immunological Genome Project (Immgen) (77). Our analysis identified several genes with specific expression in macrophages, such as *Padi4*, *Lpl*, and *Cd5l* (**Fig. 7E**). We also found high expression of *Lpl*, *Dock9*, and *Acvrl1* in macrophages and dendritic cells. As expected, genes related to immunoglobulins, namely *Igkv15-103*, *Igkv16-104*, *Igh2b*, and *Jchain*, were primarily expressed in B-cells. We also observed high expression of *Nrgn* in innate lymphocytes, while *Stfa2/3*, *Acvrl1*, *Zfyve9*, and *Mmp25* were expressed in monocytes and granulocytes, and *Igf2pb3* was mostly expressed in B-cells.

## DISCUSSION

How does maternal obesity influence future progenies to increase their risk of obesity and metabolic diseases? Recent investigations showed that *in utero* exposure to maternal obesity increases the risk of adult metabolic diseases, a phenomenon called *developmental programming* (4, 11). We postulate that such developmental programming due to maternal obesity occurs in stem cells and progenitors in a growing embryo, which can still be detected in the progenitor cells of the offsprings at a very early age when they are asymptomatic for obesity or metabolic disorders. Therefore, we performed a multi-omic analysis of bone marrow tissue that harbor various progenitor cells for immune, adipose, and mesenchymal cells that infiltrate various tissues and organs of the body.

Dysregulation of immunometabolism, the interplay between immunological and metabolic processes (78), has been reported in obesity (79), diabetes (80), and aging (81), with immunometabolic reprogramming being shown to affect the differentiation, function, and maintenance of various immune cell lineages (47, 82).

The bone marrow microenvironment is crucial in the regulation of hematopoietic stem cells. We hypothesized that maternal obesity influences the offspring bone marrow microenvironment via exposure to adverse maternal factors such as inflammation, abnormal energy metabolism, and epigenetic modifications, all of which can affect the fate and metabolism of hematopoietic stem cells. We have recently shown significant metabolic changes in the bone marrow cells of the offspring of HFD-fed mothers, namely reduced amino acid levels and alterations in energy metabolism and immune complexity (18).

Here, we conducted lipidomics analysis on bone marrow cells collected from newly- weaned offspring of mothers fed either the RD or a HFD. Surprisingly, despite increased maternal and offspring adiposity, three-week-old Off-HFD mice exhibited a significant decline in TAG levels within bone marrow cells in general and particularly in CD11b+ - enriched bone marrow cells. We have recently reported that mitochondrial respiration is increased in the bone marrow of three-week-old Off-HFD mice (18). Given that we found the reduced TAGs in bone marrow cells in Off-HFD, it is likely that lipids are being broken down and mobilized via lipolysis. This suggestion is further supported by the fact that expression of the gene Lipoprotein lipase (*Lpl*) is significantly increased in CD11b+ cells from the bone marrow of Off-HFD vs. Off-RD mice. Lipolysis can be activated by either lipid-droplet-associated or autophagy-mediated lipases (83). Autophagy-related lysosomal acid lipase (LAL) hydrolyzes both CE and TAGs to free cholesterol and free fatty acids. Our data revealed significant decreases of CE and TAG levels in Off-HFD mice; however, the exact mechanisms remain to be elucidated. Interestingly, a similar decrease in TAG levels has previously been shown in the bone marrow of patients with Gaucher’s disease, a lipid storage disorder associated with deficient activity of the lysosomal hydrolase (84).

Importantly, CD11b+ cells showed upregulation of three long-chain TAGs: 56:3; 56:4, and 58:6.The effects of long-chain TAGs on bone marrow function have previously been studied by Beau et al. (85), who reported that incubation of human bone marrow cells with long-chain TAGs strongly inhibited colony formation. In addition, accumulation of long-chain TAGs has been observed in macrophages stimulated with lipopolysaccharide and interferon-γ, indicating that synthesis of TAGs is required for inflammatory macrophage function (86). Indeed, increased fatty acids is an integral part of the inflammatory phenotype (87). At the same time, pathway analysis of differentially expressed genes in bone marrow CD11b+ cells identified macrophage alternative activation signaling as the pathway most significantly affected by maternal obesity. This raises an intriguing possibility that maternal obesity forms a unique myeloid cell phenotype characterized by lipidomic, metabolomic, and transcriptomic changes that allow proper immune function to be maintained. However, this adaptive response may have detrimental consequences in the offspring’s future life. In fact, our data revealed age-dependent deterioration of mitochondrial function in Off-HFD, with 16-week-old Off-HFD mice showing significantly reduced oxidative phosphorylation in bone marrow cells, a phenomenon not observed in Off-RD mice. Thus, what may have evolved as adaptation to an adverse maternal environment at the beginning of life may affect responses to immune challenges that appear later in life, for example cancer.

A surprising discovery from this study is the reduction in CEs found in the bone marrow cells of Off-HFD. We reason that this could reflect decreased storage CEs in lipid droplets. Previous research by Dumesnil et al. (50) has demonstrated that TAGs play critical roles in helping CEs assemble in lipid droplets, especially when there is high cholesterol intake. Therefore, we speculate that a critical concentration of TAGs is necessary to facilitate the storage of CEs in lipid droplets, which appears to be lacking in Off-HFD mice.

Prolonged changes in cholesterol availability caused by abnormally low levels of CE, similar to those we observed in the bone marrow of Off-HFD mice, may in the long run impair mitochondrial biogenesis and function and reduce oxidative phosphorylation. Loss of mitochondrial function, in turn, has been shown to reduce the cholesterol efflux to bone marrow cells and dysregulate cholesterol homeostasis, thereby activating a feedforward mechanism that results in further deterioration of mitochondrial function (88). Alternatively, reduced mitochondrial respiration in adult Off-HFD mice might be explained by long-term accumulation of 3-hydroxy-3-methylglutaric acid, a phenomenon previously linked to mitochondrial toxicity in both humans and animal models (53, 54). Ultimately, the functional consequences of impaired oxidative phosphorylation for the bone marrow cells of Off-HFD mice are currently unclear.

We also observed changes in levels of PC and PE plasmalogens in bone marrow cells from Off-HFD mice including increases in all PE plasmalogens and a decrease in several classes of PC-plasmalogens. Changes in plasmalogens were previously linked to aging, chronic inflammation, and cardiac, metabolic, and neurodegenerative diseases (89–91). Importantly, mitochondrial dysfunction has been shown to affect plasmalogens because of the high vulnerability of their vinyl-ether linkage to oxidative stress (92). The changes in plasmalogen metabolism and homeostasis have also been shown to promote mitochondrial dysfunction (93), suggesting the presence of feedforward mechanisms.

An intriguing question that remains to be addressed is the source of the perturbations in lipid metabolism that develop in bone marrow in response to maternal obesity. Bone marrow adipocytes, for example, play critical roles in maintenance of immunity, specifically in the survival of immune cells that lack the ability to uptake, store, or process fatty acids (94). This function of bone marrow adipocytes might produce the significant changes in immune complexity that we previously reported in the bone marrow of newly-weaned Off-HFD mice (18). We also found aspartate levels to be significantly upregulated in the CD11b+ bone marrow cells in Off-HFD. Interestingly, it is shown that ETC activity significantly support the aspartate production in proliferating cells (57). Aspartate is a necessary constituent for nucleotide biosynthesis as well as for protein synthesis in the cells. It will be interesting to study if increased oxidative phosphorylation seen in Off-HFD (18) is underlying the increase in aspartate levels and whether this can cause relative increase in proliferation of the myeloid progenitors in the bone marrow.

The RNA-seq analysis in this work also provided some initial insight into the bone-marrow- related programming effects, as it revealed a number of signaling pathways in CD11b+ cells to be affected by maternal obesity. Those included B-cell development and receptor signaling, consistent with our previous finding of a B-cell population uniquely represented in the bone marrow of Off-HFD mice (18). Other impacted pathways included macrophage alternative activation, the IL-7, IL-12, and IL-15 pathways, and TGFβ signaling. In Off-HFD mice, we detected increased expression of the gene encoding Apoptosis inhibitor of macrophage (AIM), also called CD5 antigen-like (CD5L), a protein that promotes autophagy and polarization of macrophages (95). Our data also revealed increased expression of the gene encoding Lipoprotein lipase (LPL), a rate-limiting enzyme in triglyceride catabolism that is expressed by macrophages (96).

Collectively, our work revealed substantial perturbations in lipid metabolism in the bone marrow of young offspring of obese mothers. Especially, we identified abnormal metabolic and transcriptomic changes in bone marrow CD11b+ cells of Off-HFD mice. Our findings raise the possibility of targeting lipid metabolism for therapeutic purposes in individuals born to obese mothers. Furthermore, understanding the nature of the factors that act within the bone marrow milieu could facilitate the prevention of immune dysfunction in the offspring of obese mothers.

## Grant Support and Aknowledgements

Financial support for this work was provided by the NIH HL170097, HL16447, HD099367, and NIDDK Mouse Metabolic Phenotyping Centers (National MMPC, RRID:SCR_008997, www.mmpc.org) under the MICROMouse Program, grants DK076169, Exploratory Research Seed Grant funding from the OHSU School of Medicine, OHSU Medical Research Foundation award (to AM); and Pacific Northwest National Laboratory (PNNL)-OHSU PMedIC project (to AM, SK, and TM). For contribution to the transcriptomics analysis, the authors acknowledge the support of the OHSU Exploratory Research Seed Grant funding, OHSU Massively Parallel Sequencing Shared Resource (MPSSR) as well as the ONPRC Bioinformatics & Biostatistics Core, which is funded in part by NIH grant OD P51 OD011092. Bone marrow cell lipidomics analysis was performed by the West Coast Metabolomics Center at UC Davis. Metabolomics and lipidomics analyses of myeloid cells were performed in the Environmental Molecular Sciences Laboratory, a U.S. DOE national scientific user facility located on the campus of PNNL in Richland, WA. Battelle operates PNNL for the DOE under contract DE- AC05-76RLO01830.

## Author Contributions

YA, EP, and TW- performed experiments with echo MRI and Seahorse analyzer, harvested bone marrow cells, and purified CD11b+ cells; TOM, CDN and SPC– conducted metabolomics and lipidomics analyses at PNNL; SPC- analyzed the lipidomics and metabolomics data; PD and RS – conducted RNA sequencing at OHSU Massively Parallel Sequencing Core and performed the bioinformatics analysis of the data; SR and SF-statistical analysis of RNA-sequencing data; BC–provided access and guidance with the echo MRI experiments; TOM, SK and AM – conceptualization and design of the study; AM - funding acquisition and data analysis; SK, TOM, and AM wrote the manuscript. All authors provided feedback and assisted with preparing the final manuscript.

## LIST OF ABBREVIATIONS

Acvrl1: Activin A receptor like type 1
BMCs: Bone marrow cells
Cd5l: CD5 Antigen-Like
CE: Cholesteryl ester
Cstdc4/5/6: Cystatin domain containing 4, 5,6
DAGs: Diacylglycerols
Dock9: Dedicator of Cytokinesis 9
HMG: 3-Hydroxy-3-methylglutaric acid (HMG)
HFD: High fat diet
Igf2bp3: Insulin Like Growth Factor 2 MRNA Binding Protein 3
Ighg2b: Immunoglobulin Heavy Constant Gamma 2b
Igkv15-103: Immunoglobulin kappa chain variable 15-103
Igkv16-104: Immunoglobulin kappa variable 16-104
IPA: Ingenuity Pathway Analysis
Jchain: Immunoglobulin J chain
Ldlrad3: Low-density lipoprotein receptor class A domain-containing protein 3
LPC: Lysophosphatidylcholine
Lpl: Lipoprotein lipase
Mmp25: Matrix metalloproteinase-25
Nrgn: Neurogranin
Off-HFD: Offspring of mothers fed a high fat diet
Off-RD: Offspring of mothers fed a regular diet
Otulinl: OTU deubiquitinase with linear linkage specificity
Padi4: Protein-arginine deiminase type-4
PNNL: Pacific Northwest National Laboratory
PC: Phosphocholine
PE: Phosphatidylethanolamines
RD: Regular diet
Stfa2/3: Stefin-2/3
TAG: Triacylglycerols
TGF β: Transforming growth factor beta
Trio: Triple functional domain protein
Zfyve9: Zinc finger FYVE domain-containing protein 9

---

**Supplemental Figure 1.**
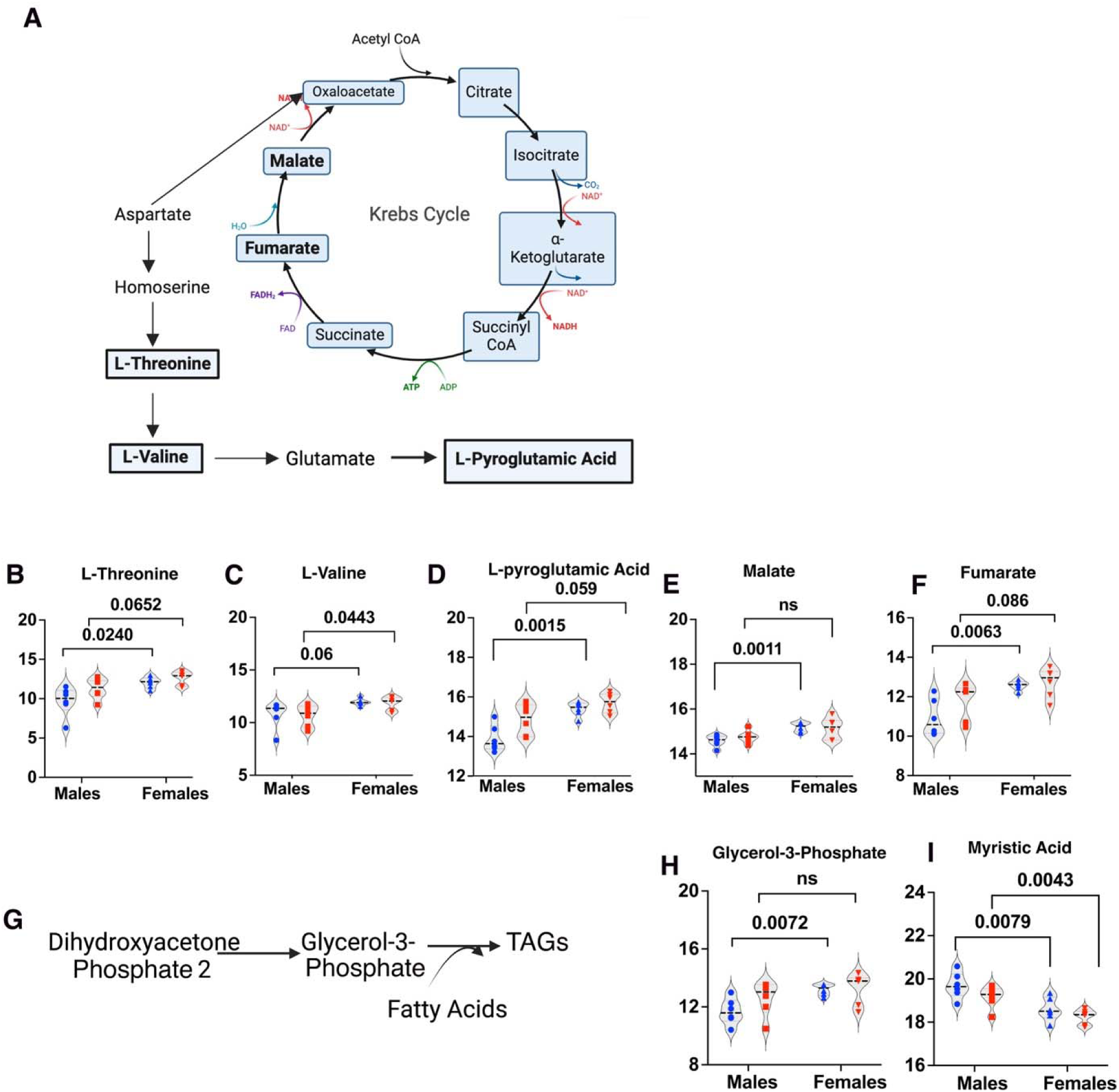
Sex-dependent differences in metabolite levels in bone-marrow- derived CD11b+ myeloid cells. **A**, Schematic of amino acid synthesis from L-aspartate and in the Krebs cycle. **B-I**, Sex-dependent changes in metabolite levels according to maternal diet group. Affected amino acids: **B**, L-threonine; **C**, L-valine; and **D**, L-pyroglutamic acid. Altered Krebs cycle metabolites: **E**, malic acid and **F**, fumaric acid. **G**, Schematic of the triglyceride synthesis pathway. Affected triglyceride metabolites: **H**, glycerol-3-phosphate and **I**, myristic acid. *P*-values are shown; ns, not significant. Illustrations in A and G created with BioRender.com.

